# Microbiome-TP53 Gene Interaction in Human Lung Cancer

**DOI:** 10.1101/273524

**Authors:** K. Leigh Greathouse, James R. White, Ashely J. Vargas, Valery V. Bliskovsky, Jessica A. Beck, Natalia von Muhlinen, Eric C. Polley, Elise D. Bowman, Mohammed A. Khan, Ana I. Robles, Tomer Cooks, Bríd M. Ryan, Amiran H. Dzutsev, Giorgio Trinchieri, Marbin A. Pineda, Sven Bilke, Paul S. Meltzer, Alexis N. Hokenstad, Tricia M. Stickrod, Marina R. Walther-Antonio, Joshua P. Earl, Joshua C. Mell, Jaroslaw E. Krol, Sergey V. Balashov, Archana S. Bhat, Garth D. Ehrlich, Alex Valm, Clayton Deming, Sean Conlan, Julia Oh, Julie A. Segre, Curtis C. Harris

## Abstract

**Background:** Lung cancer is the leading cancer diagnosis worldwide and the number one cause of cancer deaths. Exposure to cigarette smoke, the primary risk factor in lung cancer, reduces epithelial barrier integrity and increases susceptibility to infections. Herein, we hypothesized that somatic mutations together with cigarette smoke generate a dysbiotic microbiota that is associated with lung carcinogenesis. Using lung tissue from controls (n=33) and cancer cases (n=143), we conducted 16S rRNA bacterial gene sequencing, with RNA-seq data from lung cancer cases in The Cancer Genome Atlas (n=1112) serving as the validation cohort.

**Results:** Overall, we demonstrate a lower alpha diversity in normal lung as compared to non-tumor adjacent or tumor tissue. In squamous cell carcinoma (SCC) specifically, a separate group of taxa were identified, in which *Acidovorax* was enriched in smokers (*P* =0.0013). *Acidovorax temporans* was identified by fluorescent *in situ* hybridization within tumor sections, and confirmed by two separate 16S rRNA strategies. Further, these taxa, including *Acidovorax*, exhibited higher abundance among the subset of SCC cases with *TP53* mutations, an association not seen in adenocarcinomas (AD).

**Conclusions:** The results of this comprehensive study show both a microbiome-gene and microbiome-exposure interactions in SCC lung cancer tissue. Specifically, tumors harboring *TP53* mutations, which can damage epithelial function, have a unique bacterial consortia which is higher in relative abundance in smoking-associated SCC. Given the significant need for clinical diagnostic tools in lung cancer, this study may provide novel biomarkers for early detection.

## Background

Lung cancer is the leading cancer diagnosis worldwide (1.8 million/year) and has a higher mortality than that of the next top three cancers combined (158,080 vs 115,760 deaths) [1]. Unfortunately, lung cancer survival remains poor and has shown minimal improvement over the past five decades, owing to diagnosis at advanced stage and resistance to standard chemotherapy [2]. While we have made significant strides with targeted receptor therapy and immunotherapy, biomarkers with higher specificity would improve diagnosis and treatment for these individuals.

Epidemiological evidence indicates an association between repeated antibiotic exposure and increased lung cancer risk, however, the contribution of the lung microbiome to lung cancer is unknown [3]. The first line of defense against inhaled environmental insults, including tobacco smoke and infection, is the respiratory epithelium. Until recently, healthy lungs were regarded as essentially sterile; however, studies now illustrate the presence of a lung microbiota [4], the community of microscopic organisms living within the host lung, which is altered in respiratory diseases including asthma, chronic obstructive pulmonary disease (COPD) and cystic fibrosis [5]. Disruption of the epithelium by tobacco smoke can be a primary cause of inflammatory pathology, which is seen in both COPD and lung cancer. Dysbiosis has been observed in both humans and model systems of COPD and cystic fibrosis [6, 7]. In COPD patients and *in vitro*, cigarette smoke has been shown to reduce epithelial integrity and cell-cell contact, which can increase susceptibility to respiratory pathogens or other environmental pollutants [8]. Disturbances in the microbiome, from cigarette smoke, epithelial damage or gene mutations, can allow pathogenic species to dominate the community or increase virulence of other normally commensal microbes. Evidence of this has been demonstrated in patients with cystic fibrosis who have more virulent forms of *P. aeruginosa* [9]. These inflammatory associated events have been proposed to lead to an increased risk or progression of diseases, including lung cancer.

Several bacteria are associated with chronic inflammation and subsequent increased risk of lung and colon cancer, including *Mycobacterium tuberculosis* (lung cancer) [10] *Bacteroides fragilis* and *Fusobacterium nucleatum* (colon cancer) [11]. Recent microbiome studies in colon cancer have demonstrated a contribution of bacteria to carcinogenesis. Specifically, *F. nucleatum,* a bacterium commonly isolated from patients with inflammatory bowel disease, may be a risk factor for colon cancer [11, 12]. The more virulent strains of *F. nucleatum* affect colon cancer progression in animal models and increase tumor multiplicity [13] by various mechanisms including favoring the infiltration of tumor-promoting myeloid cells to create a pro-inflammatory environment [14]. Colorectal carcinomas associated with high abundance of fecal *F. nucleatum* were found to have the highest number of somatic mutations, suggesting that these mutations create a pathogen-friendly environment [15]. Similarly, *B. fragilis* can secrete endotoxins that cause DNA damage leading to mutations and colon cancer initiation [16]. Furthermore, the loss of the oncogenic protein p53 in enterocytes impairs the epithelial barrier and allows infiltration of bacteria resulting in inflammatory signaling (NF-κB), which is required for tumor progression [17]. The tumor suppressor gene *TP53* is the most commonly mutated gene in lung cancer [18], with certain missense mutations showing gain of oncogenic function [19]; however, the relationship between *TP53* and microbiota in lung cancer remains unknown. Herein, we hypothesize that somatic mutations together with alterations in the lung microbiome create an inflammatory environment, which participate in lung carcinogenesis.

## Results

To investigate the lung mucosal-associated microbial alterations in the etiology of lung cancer, we analyzed samples from the NCI-MD case-control study (n=143 tumor and n=144 non-tumor adjacent tissues) and lung cancer samples from The Cancer Genome Atlas (n=1112 tumor and non-tumor adjacent RNA-seq data from tissues) for validation. In addition, we used the clinical information from these two sample populations to control for confounders in lung cancer risk and progression (age, gender, smoking, race, family and medical history and co-morbidities), as well as factors that are known to alter the human microbiome (antibiotics and neo-adjuvant therapy). Given the paucity of healthy lung tissue available for study, we utilized two separate tissue biorepositories. Non-cancerous lung tissue was obtained by lung biopsy from individuals with benign lung nodules without cancer or non-cancer lung from immediate autopsy [20], which was used as a referent control (see Additional file 1: Table 1). Given the high potential for contamination in low-biomass samples, such as the lung, we took several measures to address this issue controlling for contamination points in the collection process. In order to remove possible contaminants from our analysis, we first performed a threshold analysis similar to a previous study [21], wherein we plotted the mean percent abundance across experimental samples versus negative control samples and removed those that were ≥ 5% in both experimental and negative control samples (see Additional file 1: Figure S1). Additionally, we conducted hierarchal clustering of negative controls, non-tumor and tumor samples independently in order to visualize and identify the strongest sources of contamination (see Additional file 1: Figure S2). The combination of these analyses resulted in initial removal of the genera *Halomonas, Herbaspririllium, Shewanella, Propionibacterium* and *Variovorax*.

To identify the microbial communities present in each tissue type, we sequenced the V3-5 16S rRNA bacterial gene using the Illumina MiSeq platform. After quality filtering and contaminant removal, 34 million quality sequences were retained for OTU clustering and downstream analysis (see Additional file 1: Table 2).

To enable us to validate findings from our NCI-MD 16S rRNA gene sequencing analysis we took advantage of the TCGA lung cancer database. Using the unmapped RNA-seq reads from these samples (N=1112 and n=106 paired tumor/non-tumor), we analyzed with our metagenomics analysis pipeline. After removal of all human reads, we took the remaining non-human reads and used three separate tools, MetaPhlAn, Kraken and PathoScope, to assign reads to taxonomy, including bacteria, virus, and fungi. Due to the highly curated database of PathoScope, we were able obtain to species and in some cases strain-level putative identification of RNA-seq reads. For this reason, and due to its rigorous validation in other studies [22], we used these data as our validation data set. Unfortunately, given that all patients in this database had lung cancer, we could not validate our microbial findings in non-diseased lung tissue in the TCGA data set. Given that this was one of the first times TCGA was used to completely profile the microbiota of lung cancer, we asked how similar the 16S rRNA gene sequencing and RNA-seq microbial communities were at the phylum and genus levels. Using an overall threshold of 0.01% of genus level abundance, we identified 236 overlapping genera out of 520 total genera in the 16S rRNA gene sequencing data and 609 total genera in the RNA-seq data (see Additional file 1: Figure S3).

### Bacterial profile of the lung cancer microbiome is dominated by Proteobacteria and validated in a separate lung cancer data set

We know from previous microbial studies of lung disease that bacterial composition shifts occur compared to normal non-diseased lungs [23] and associated with disease severity [24], however these compositional changes have not been examined in lung cancer. In order to identify the microbial changes associated with lung cancer, we first examined the ecological diversity within samples (alpha diversity) and between samples (beta diversity) of non-cancerous (immediate autopsy and hospital biopsy) tissues, non-tumor adjacent (NT) and tumor (T) tissues from 16S rRNA gene sequencing. At the phylum level, we observed increases in Proteobacteria (Kruskal-Wallis *P=*0.0002) and decreases in Firmicutes (Kruskal-Wallis *P=*0.04) in lung tissue hospital biopsies, as well as, in tumor and associated non-tumor tissues from the NCI-MD study compared with non-cancer population control lung tissues, as has been seen in COPD[25] (Figure 1A). We also observed a similar increase in Proteobacteria (Mann-Whitney *P=*0.02) between non-tumor lung tissue and lung cancer in the TCGA study, indicating that this is recurrent phenomenon in lung cancer (Figure 1A). However, the lack of similarity between the NCI-MD and TCGA non-tumor samples may be attributed to the TCGA data being derived from multiple sample populations in the U.S., differences in sample prep and in sequencing platforms, as illustrated by Meisel et al. [26].

**Figure 1.**
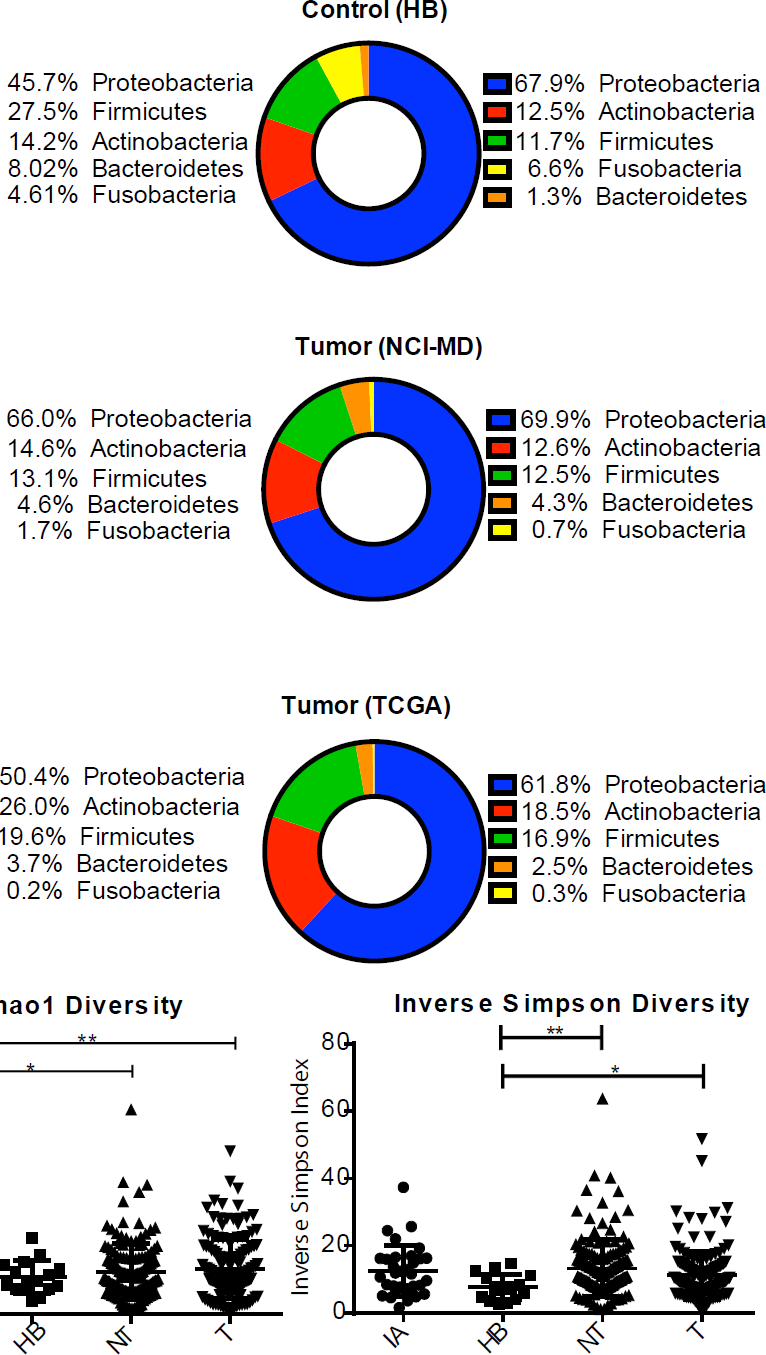
The bacterial profile and diversity of the lung microbiome in non-diseased and cancerous tissues. A) 16S rRNA gene sequences from non-diseased lung (ImA or HB; top), non-tumor adjacent (NT) and tumor (T) assigned to OTUs or proportional abundance of metatranscriptomic sequences (TCGA; bottom) at the phylum level showing the most dominant taxa for each tissue type. B) Alpha diversity between non-diseased lung tissue (ImA and HB) non-tumor adjacent (NT) and tumors from 16S rRNA gene sequencing using Chao1 (richness) or inverse Simpson index. *(*P*<0.05), **(*P*<0.01). Test of significance is Mann-Whitney. ImA - immediate autopsy; HB – hospital biopsy.

To identify ecological diversity changes associated with lung cancer, we next examined the richness (Chao1) and diversity (Inverse Simpson) of the microbiome within samples (alpha diversity) of non-disease (immediate autopsy and hospital biopsy) lung tissues, non-tumor adjacent and tumor tissues from 16S rRNA gene sequencing (NCI-MD study). Specifically, Chao1 measurement demonstrated a significant increase in both tumor and non-tumor tissue richness as compared to immediate autopsy control tissue samples (Figure 1B). Similarly, using the Inverse Simpson index, which measures number (richness) and abundance (evenness) of species, we observed a significant increase in alpha diversity in both tumor and non-tumors as compared to hospital biopsy control tissues (Figure 1B), similar to studies of severe COPD [27] indicating that microbial diversity of lung cancer tissues are altered from its non-diseased state. However, we did not see any significant changes in alpha diversity by smoking status (never, former or current) nor correlation with time since quitting smoking (see Additional file 1: Figure S3), in cancer-free or lung cancer tissues as has been demonstrated in other lung microbiome studies [28, 29]. We also asked whether there were differences between microbial communities using beta diversity (Bray Curtis). Within the NCI-MD study we observed significant differences in beta diversity between all tissue types (PERMANOVA *F*=2.90, *P=*0.001), tumor and non-tumor (PERMANOVA *F*=2.94, *P=*0.001) and AD v SCC (PERMANOVA *F*=2.27, *P=*0.005), with tumor vs. non-tumor having the largest among-group distance denoted by the higher *F* value (data not shown). Similarly, we observed significant difference in beta diversity between tumor and non-tumor (PERMANOVA *F*=3.63, *P=*0.001) and AD v SCC (PERMANOVA *F*=27.19, *P=*0.001) in addition to lung resection location (upper, lower, left, right) (PERMANOVA *F*=1.36, *P=*0.061) (data not shown). Together, these data illustrate a trend of increasing diversity and richness associated with lung cancer.

### A distinct group of taxa are enriched with squamous cell carcinoma (SCC) with *Acidovorax* more abundant in smokers

The two most common types of non-small cell lung cancer are SCC and AD, arising centrally from the cells lining the bronchi and from peripheral airways, respectively. Previous studies report that the microbial community differs between the bronchi and lower lungs in COPD [6]. This phenomenon of anatomic-specific microbial variation was also apparent in the abundance of genera between bronchial and SCC tumors from the upper lungs with higher abundance of *Acidovorax* in comparison to AD tumors (see Additional file 1: Figure S4). Further, the taxonomic distribution in AD tumors appears more similar to the taxonomic abundance in COPD, which is generally dominated by *Pseudomonas* [6]. Given this distinction, we controlled for this potential confounder of lung location in subsequent analyses. This led us to investigate the specific taxonomic pattern further and ask if there was a specific microbial consortia that is enriched in SCC or AD tumor tissue. In the NCI-MD study we identified 32 genera that were differentially abundant in SCC (n=47) vs AD (n=67) tumors (Student’s t-test; MW *P*<0.05), 9 of which were significant after multiple testing correction (FDR) (*Acidovorax, Brevundimonas, Comamonas, Tepidimonas, Rhodoferax, Klebsiella, Leptothrix, Polaromonas, Anaerococcus*) (Figure 2A). We also validated these same observations in the TCGA data set (AD=485, SCC=489) (Mann-Whitney FDR corrected p-value<0.05) (Figure 2B). To control for potential confounders of this association, including age, gender, race, smoking, anatomical location and stage, we conducted adjusted logistic regression analysis in the NCI-MD study for each taxa separately and confirmed 6/9 of these genera were significantly associated with increased odds of being SCC as compared to AD lung cancer (Figure 2C and see Additional file 1: Table 5). Though we had reduced power, we asked whether the time since quitting smoking would change this association, and found that *Acidovorax, Klebsiella, Tepidimonas, Rhodoferax,* and *Anaerococcus* remained significant. When we examined the larger TCGA data set, we also found significantly increased odds of being SCC as compared with AD among 4/9 (*Acidovorax, Klebsiella, Rhodoferax, Anaerococcus*) of the same genera in adjusted models (FDR corrected *P*<0.05) (Figure 2D and see Additional file 1: Table 6). This association also remained significant after adjusting for pack years and time since quitting smoking. Together these data, validated in two separate cohorts, demonstrate that a specific community of taxa is more abundant in SCC as compared with AD lung cancer tissue, and are capable of distinguishing between AD and SCC tumors from individuals with similar exposure to cigarette smoke. However, whether this is a cause or consequence of the development of SCC cancer remains unknown.

**Figure 2.**
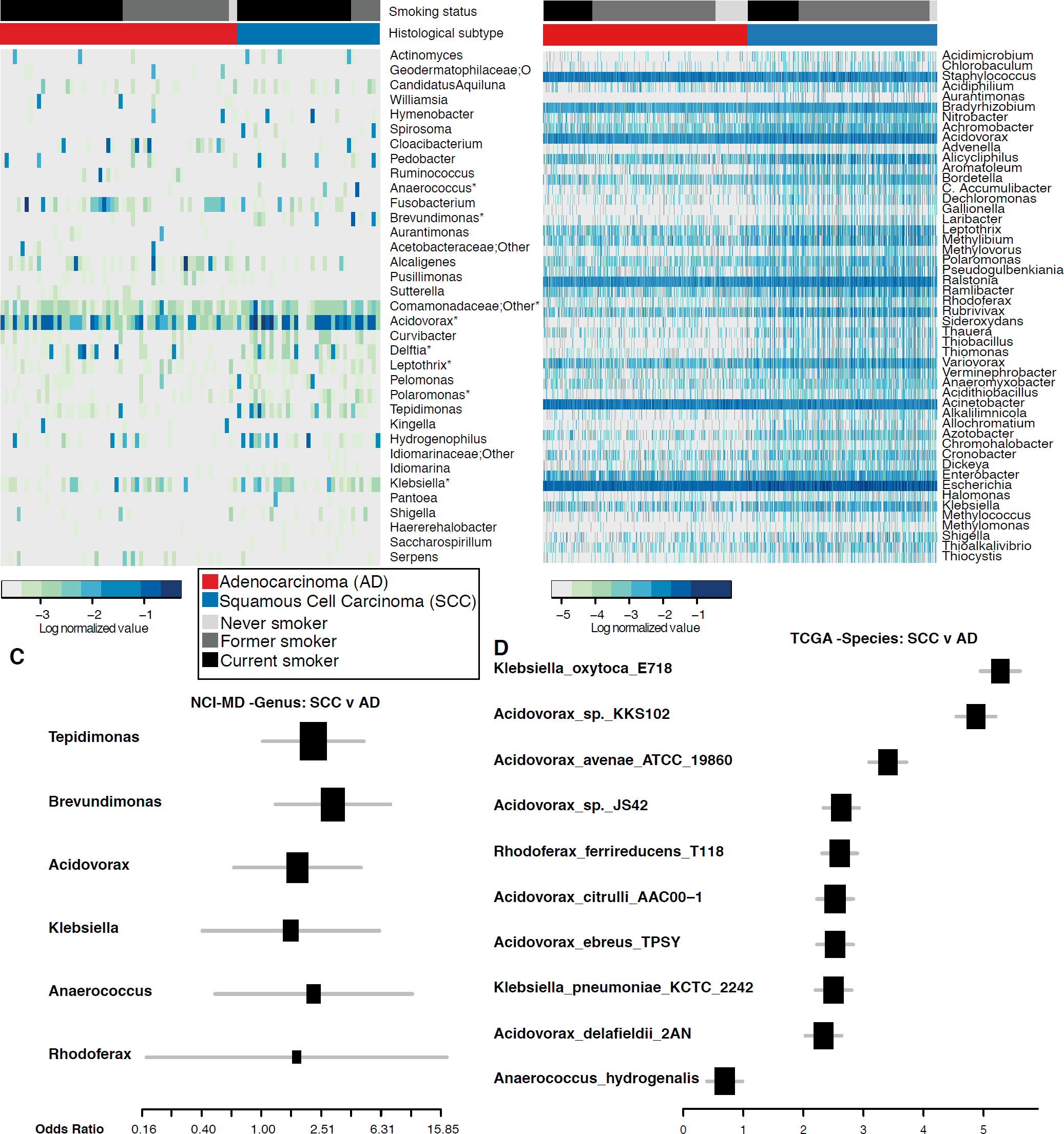
Taxonomic consortia differentiating smoking status and histological subtype of lung cancer. A) Heat maps showing top differentially abundant genera (NCI-MD) (Mann-Whitney p-value<0.05; **\*** overlapping between NCI-MD and TCGA) between AD and SCC lung cancer tissue sorted by histological subtype and smoking status. B) Heat map showing genera (TCGA) that that are differentially abundant between AD and SCC (Mann-Whitney FDR corrected *P*<0.05), sorted by histological subtype and smoking. C) Forest plot of odds ratios for genera in NCI-MD data set that are significantly associated with SCC compared to AD in tumors (adjusted odds ratio *P*<0.05). D) Forest plot of odds ratios for species in TCGA data set that are significantly associated with SCC versus AD in tumors (adjusted odds ratio FDR corrected *P*<0.05).

Both SCC and AD lung cancers are associated with smoking, however, the association between smoking and SCC is stronger [30], which lead us to ask whether any of the SCC-enriched taxa were also associated with smoking. We stratified the tumor samples into never smokers (n=7) or ever-smokers (current (n=70) and former smokers (n=40)) using linear discriminant analysis (LEfSe) to identify smoking-associated microbial biomarkers in SCC tumors. We identified 6 genera that were able to distinguish ever (former and current) vs non-smokers in our NCI-MD study (*Acidovorax, Ruminococcus, Oscillospira, Duganella, Ensifer, Rhizobium)* (see Additional file 1: Figure S4C). Specifically, *Acidovorax* was more abundant in former and current smokers as compared with never smokers (Kruskal-Wallis p-value <0.05) (Figure 3A), with a similar trend observed in TCGA data set (n_never_=120, n_former_=551, n_current_=217) (Kruskal-Wallis *P*=0.27; ANOVA *P*=0.02). We did not, however, observe any correlation between *Acidovorax* abundance and smoking time cessation. Interestingly, the relative abundance of *Acidovorax* and *Klebsiella* were higher in former and current smokers when we stratified by histological subtype in both NCI-MD and TCGA data set (Figure 3B, see Additional file 1: Figure S5), indicating not only are there bacteria which have a higher relative abundance in tumors from individuals who smoke, but SCC tumors from smokers have even greater relative abundance of these bacteria. We also demonstrated the presence of this bacterium in lung tumors using FISH (Figure 3C-D and see Additional file 1: Figure S6), and using PacBio sequencing, which identified the species as *A. temperans* (see Additional file 1: Table 4). We did not find any significant associations between pack years or time since quitting smoking and the abundance of these taxa in either study among SCC tumors in either study.

**Figure 3.**
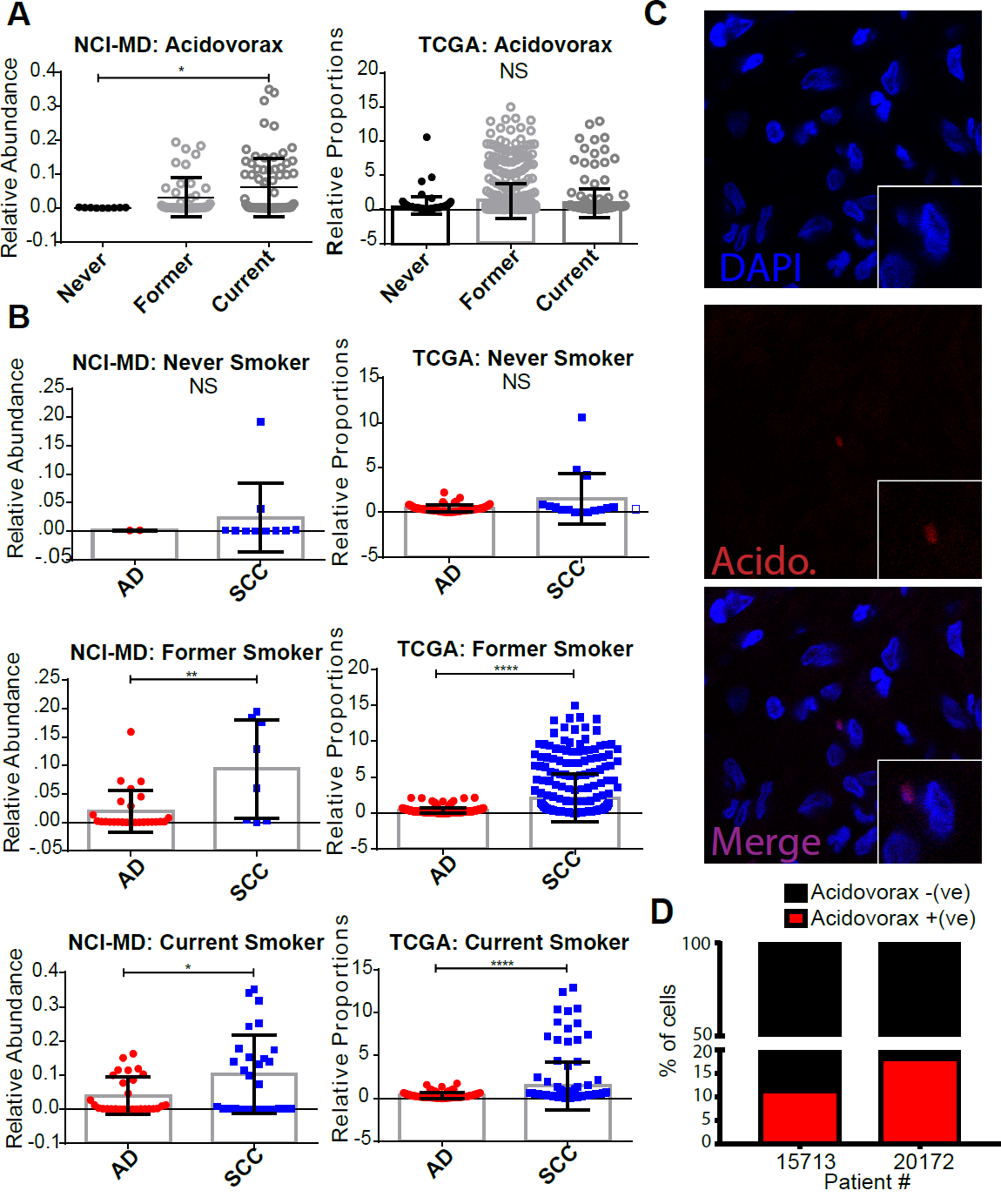
Relative abundance of *Acidovorax* stratified by smoking status and histological subtype. A) Relative abundance of *Acidovorax* stratified by smoking status in NCI-MD (Left) and TCGA (Right) data sets. B) Relative abundance of *Acidovorax* in Never, Former and Current smokers stratified by histological subtype in the NCI-MD (Left) and TCGA (Right) data sets. C) Representative FISH images of tumor tissue sections using fluorescent probe specific to *Acidovorax*. D) Quantification of *Acidovorax* probe reactivity (10 fields; at least 300 cells counted) showing percentage (%) of cells with perinuclear probe reactivity from two lung cancer cases (15713 – SCC/current smoker; 20172 – SCC/former smoker). *(*P*<0.05), **(*P*<0.01), ****(*P*<0.0001). Test of significance are Mann-Whitney or Kruskal-Wallis and Dunn’s multiple comparisons test. NS = non-significant.

### *TP53* mutations are associated with enrichment of SSC enriched taxa

The most prevalent somatic mutation in SCC lung tumors is in the gene *TP53* [31]. Previous studies demonstrate that mutations in *TP53*, specifically in colon cancer, lead to disruption of the epithelial barrier allowing the infiltration of tumor-foraging bacteria and resulting in disease progression [17]. Given that *TP53* mutations are found in 75-80% of SCC tumors, we hypothesized that these SCC-associated taxa may be more abundant in tumors with *TP53* mutations, owing to the loss of the epithelial barrier function in these tumors. To address this question, we investigated the association between *TP53* mutations in both NCI-MD (n=107) and TCGA (n=409) data sets using either *TP53* specific sequencing (MiSeq) or the published *TP53* mutation analysis data from TCGA [31]. We first analyzed all tumors in the NCI-MD study regardless of histology and identified a group of taxa that were more abundant in tumors with *TP53* mutations (Figure 4A). To have greater power, we performed the same analysis in the TCGA data set and observed a significant increase in these same taxa (MW FDR corrected *P*<0.05) (Figure 4B). When analyzing only SCC tumors (n=46), this signature became stronger in tumors with *TP53* mutations in both data sets, specifically among the SCC-associated taxa previously identified (Figure 4C-D). In the NCI-MD study we found that 5/9 of the genera (*Acidovorax, Klebsiella, Rhodoferax, Comamonas,* and *Polarmonas*) that differentiated SCC from AD were also more abundant in the tumors harboring *TP53* mutations, though not statistically significant (Figure 4C). In the TCGA data set, the fold change in all 5 SCC-associated genera were significantly higher in SCC tumors (n=177) with *TP53* mutations (MW corrected FDR <0.01; Figure 4D). Furthermore, using these same SCC-associated taxa we observed no pattern of association in AD tumors with *TP53* mutations indicating this signature was specific to SCC with *TP53* mutations (see Additional file 1: Figure S7A-B). Overall, these data are consistent with the hypothesis that mutations in *TP53* are associated with the enrichment of a microbial consortia that are highly represented in SCC tumors.

**Figure 4.**
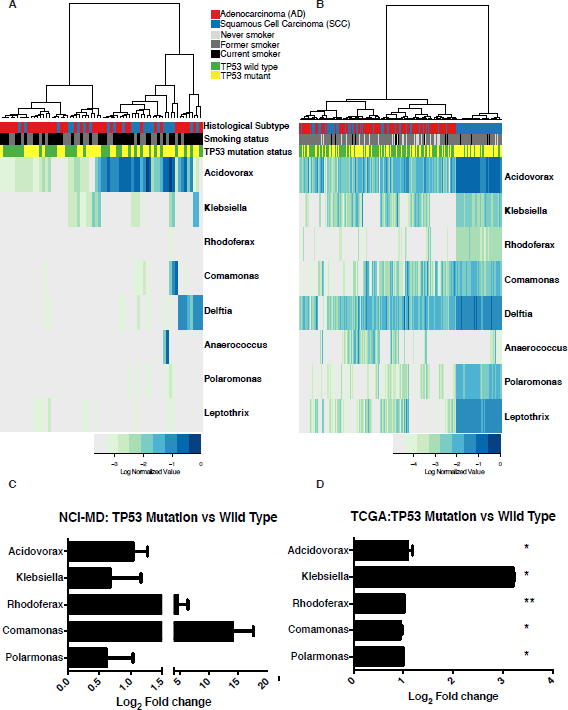
Mutations in *TP53* associated with abundance of taxonomic signature specific to squamous cell lung tumors. A) Heat map of genus-level abundance in NCI-MD data colored by mutation status, *TP53* wild type or mutated, smoking and histological subtype in all lung tumor samples. B) Heat map of genus-level abundance from TCGA data in all tumors colored by mutation status, *TP53* wild type or mutated, smoking and histological subtype. C-D) Fold change in mean abundance of SCC-associated taxa in NCI-MD or TCGA tissues comparing *TP53* mutated to wild type. Tests of significance are Mann-Whitney. Fold change among all taxa in D) are significant after FDR correction <0.01. (NCI-MD; SCC_wt_=11, SCC_mut_=35 and TCGA; SCC_wt_=59, SCC_mut_=118)

## Conclusions

Gene-environment interactions have been identified as contributors to cancer incidence [32], however little is known about gene-microbiome interactions in carcinogenesis. We demonstrate, a gene-microbiome association in human lung cancer, as well as, histological evidence of a smoking-associated bacterium, *Acidovorax*. Herein, we identify a microbial consortia that is associated with a histological subtype of lung cancer, SCC, which is further enriched in tumors with mutations in *TP53*. Given the strong association between smoking and development of SCC, it follows that a subgroup of this SCC consortium would also be found in smoking-associated SCC. We validate this assumption finding *Acidovorax spp.* more abundant in SCC tumors harboring *TP53* mutations, and confirmed the presence of this genus histologically. These results suggest that smoking may provide an environment conducive to the growth of *Acidovorax spp*. and similar species, which can flourish in nutrient depleted environments, like that of the lung. More important these bacteria are capable of using and degrading environmental toxicants, like those found in tobacco smoke [33]. Alteration of the microenvironment could allow these species to become tumor-foraging bacteria once the epithelial barrier defense is lost as a consequence of mutations in *TP53* and malignant transformation. The counterfactual is also possible. In other words, overgrowth of *Acidovorax spp.*, as the result of smoking and epithelial barrier damage, may induce mutations by activation of carcinogens, and propagation of mutated epithelial cells. Whether these species are simply opportunists or contribute to lung cancer progression, should be the subject of future investigations. Collectively, these observations indicate that a state of dysbiosis exists in lung cancer. The hypothesis generated is that epithelial cells in the lung exposed to tobacco smoke and/or mutations in *TP53* are invaded by species that take advantage of this new microenvironment. Notably, individuals harboring mutations in *TP53* with stage I SCC also have poorer prognosis [34], thus it will be important to determine if any of the species enriched in SCC are functionally related to reduced survival or simply biomarkers of a diminished mucosal barrier function. Future studies could test the hypothesis using germ-free mouse models of lung cancer inoculated with SCC-associated spp. prior to or after lung tumors are present.

Our study indicates that smoking is associated with alterations in relative abundance of species in SCC tumors. The number one risk factor for lung cancer is tobacco exposure and is a known factor in chronic lung inflammation. Tobacco and cigarette smoke contain bacterial products (i.e. LPS) that can cause inflammation, impaired barrier function and potentially alter the microbiome to influence lung carcinogenesis [8, 35, 36]. Additionally, tobacco leaves harbor both mold and potentially pathogenic bacteria that can be transferred in a viable form into the respiratory tract on tobacco flakes inhaled in mainstream smoke [35, 36]. Further, biologically significant quantities of bacteria are microaspirated daily in healthy individuals [37], and thus is possible for these species to accumulate in a pathogen-friendly environment. Given that *Acidovorax spp.*, which have been identified in 2 common brands of cigarettes [38], have the capacity to metabolize multiple organic pollutants, like those found in cigarette smoke, it is plausible that smoking creates an environment that allows this bacterium to outcompete other species for resources and thus survival [39]. Further, these bacteria are capable of metabolizing polycyclic aromatic hydrocarbons, which are tobacco smoke carcinogens that can cause DNA adducts, suggesting they may actually contribute to lung carcinogenesis.

Oral and nasal microbiome differences have been observed between smokers and non-smokers [40, 41]. Whether smoking alters the lung *tissue* microbiome; however, is still not well understood, especially in the context of disease. From a large study of the naso- and oropharynx, significant differences in microbial communities were identified between smokers and non-smokers [42]. Additionally, in a study of non-malignant lung tissue (n=152), they observed a significant increase in alpha diversity with higher number of pack years of smoking [43]. While they identified *Acidovorax, Anaerococcus* and *Comamonas* in smokers, these taxa did not differentiate smokers and non-smokers in a *healthy* population. However, in a recent study of non-malignant lung tissue, which compared tissue to isolated extracellular vesicles (EVs) from tissues, the greater diversity was identified specifically in EVs, with a greater abundance of *Acidovorax* specifically found in the EVs of smokers, indicating a possible factor in differential findings [44]. These data indicate that smoking alone may be insufficient to alter the microbial population in a healthy population. However, smoking has been shown to suppress the immune system and induce epithelial barrier dysfunction [45]. These factors may allow taxa direct access to epithelial cells where microbial toxins or reactive oxygen/nitrogen from the aforementioned species to directly or indirectly encourage malignant transformation of the lung epithelium via DNA damage and mutations in *TP53* [46-48].

Several cancers are caused by bacteria and viruses, including cervical cancer (HPV), liver cancer (HBV), gastric cancer (*H. pylori* and potentially *B. fragilis)*. However, very few microbes have been identified as carcinogenic. In contrast, microbes such as *H. pylori* and *E. coli* can promote carcinogenesis through modulation of inflammation, reactive oxygen species, metabolism and protein degradation. Specifically, several species have been shown to modulate the tumor-suppressor p53 at both the protein and DNA level [49]. Mutation in *TP53* is a common hallmark of cancer, and in SCC of the lung is found to be mutated in 70-80% of tumors [31, 50]. Mutation of *TP53* allows the tumor cells to evade immune surveillance, cell cycle arrest and death, as well as, disrupt endothelial cell tight junctions[51]. Colorectal carcinomas associated with high abundance of fecal *F. nucleatum* were found to have the highest number of somatic mutations, suggesting that these mutations create a pathogen-friendly environment [11, 15]. Furthermore, the loss of p53 in enterocytes impairs the epithelial barrier and allows infiltration of bacteria resulting in NF-κB signaling, which was required for tumor progression [17]. This evidence suggests that SCC tumors with *TP53* mutations could have poor epithelial barrier function, thus allowing tumor foraging bacteria, like those identified in our study, to become more abundant in tumors with *TP53* mutations. The counterfactual is also possible. Similar to the *B. fragilis* toxin ETBF, which is genotoxic and initiates colon carcinogenesis in animal models [52], one or more of the tumor-associated species may induce *TP53* mutations. Whether any of these bacteria are promoting SCC tumorigenesis or inducing mutations in *TP53* is currently under investigation.

With the majority of lung cancer being diagnosed at late stage, the recent advancement in the treatment of late stage (III/IV) lung cancer with immune checkpoint inhibitors targeting PD-1, nivolumab, has resulted in a 40% reduced risk of death as compared to standard chemotherapy [53]. The response rate, however, is still not complete for these patients. Important insights into understanding the differential response rates of this new immunotherapy has suggested the composition of the lung microbiome prior to therapy as a key player in therapeutic effectiveness [54]. Given our results demonstrating alterations in the microbial composition in lung cancer that are histology and mutation specific, future studies should address whether the lung or nasal microbiome composition improves the stratification of patients who would be most responsive to immunotherapy. This suggestion is supported by recent animal studies demonstrating the contribution of the gut microbiome to the effectiveness of immunotherapy [55].

The strength of our findings include the large number of individuals sampled in this study, use of two separate sample populations, two sets of control populations, two separate sequencing methodologies (MiSeq and PacBio) and microscopic validation (FISH) of the species in lung tumor tissue. We have also been diligent in assessing the possibility of contaminating taxa being an artifact of sample collection or sample processing by sequencing across 2 different platforms, and microscopy, although we cannot rule that possibility out entirely. While we were able to control for antibiotic exposure in the NCI-MD study, we acknowledge a limitation of the validation study is the inability to control for antibiotic exposure in the TCGA dataset, as well as, significant differences in clinical features between the cancer cases and controls, which could be confounders. However, in a recent study of the microbiome of endoscopic gastric biopsies, confirmation of multiple shared bacteria in clinical samples, specifically *H. pylori*, was demonstrated using the TCGA RNA-seq data with methods similar to those presented in our study [56]. With these results, we foresee a new avenue for mechanistic studies to address the role of microbe-host relationship in lung cancer inflammation, response to therapy, and microbial engineering for drug delivery.

## Methods

### Sample Populations and data sets

Samples used for DNA extraction, PCR and sequencing were obtained from the ongoing NCI-MD study (7 hospitals participating in the greater Baltimore, MD area recruited from 1999 to 2012), as described previously [57], from which 398 lung cancer cases were obtained, and included both tumor and non-tumor adjacent, with 121 matched pairs. The final sample set used for analysis after sequencing, which contained 106 matched pairs after quality control, is found in See Additional file 1: Table 1. Lung tumors and paired non-tumor adjacent samples from the NCI-MD study were obtained at the time of surgery, from which a section of tumor and non-involved adjacent lung tissue were flash frozen and stored at -80 degrees. At the time of study entry, a detailed patient interview was conducted to obtain basic clinical information in addition to previous cancers, neo-adjuvant therapies, current medications, family history of cancer, smoking history, education level and financial status. Staging was assigned using the Cancer Staging Manual of the American Joint Committee on Cancer (AJCC) 7th edition. Pre-operative antibiotics were administered for those cases recruited after 2008 and any antibiotic oral medication use was used as a co-variate for all statistical analysis in model testing, however these data were not available for immediate autopsy (ImA) non-cancer samples. Controls representing non-cancerous tissue were obtained from the Lung Cancer Biorepository Research Network (n=16; hospital controls). Theses samples were obtained as frozen lung specimens from individuals who had a previous positive nodule identified by PET scan, and subsequently underwent tissue biopsy, which was ruled benign. Clinical information included those listed above as well as smoking history, antibiotic usage (Y/N), and disease diagnosis. Two cases had emphysema at the time of biopsy and were not used in the analyses. Immediate autopsy (ImA) samples obtained from the University of Maryland (UMD) hospital, which is part of the NCI-MD study population (n=41; population controls) (See Additional file 1: Table 1). Lung tissue from ImA was received frozen from the UMD biorepository and served as the population controls for non-cancer lung tissue. Briefly, samples from ImA were obtained within minutes (<30 min.) after death and put on ice for dissection or frozen to -80 degrees. All ImA subjects underwent extensive autopsy and were determined to be cancer free. Demographic information included age, gender, race and cause of death only. Non-smokers in the NCI-MD study were categorized as having smoked <100 cigarettes or fewer than 5 packs over a lifetime, whereas smokers were categorized as current smokers or formers smokers, which had quit for > 6 months. Sequences derived from RNA-seq of lung tumor (n=1006) or non-tumor adjacent tissue (n=106) were obtained from The Cancer Genome Atlas (N=1112) for validation of the NCI-MD study16S rRNA gene sequencing analysis and results. Due to the fact that all RNA-seq data in TCGA were obtained using poly-A capture, any microbial data from this analysis will necessarily be biased. For this reason, we only used these data as validation of results first identified in our 16S rRNA gene sequencing analysis. Public data, including all clinical patient information (See Additional file 1: Table 1), was downloaded from the Data Matrix on the TCGA website, https://tcga-data.nci.nih.gov/tcga/dataAccessMatrix.htm. The raw data in the form of BAM and FastQ files were download from a secure server at CGHUB, and access was applied for and approved for raw data downloads by University of California Santa Cruz, https://cghub.ucsc.edu/. The files were downloaded and stored in archived format and subsequently un-archived for analysis. The results shown here are in whole or part based upon data generated by the TCGA Research Network: http://cancergenome.nih.gov/

### DNA extraction and 16S rRNA gene sequencing

DNA from lung cancer and control lung tissues was isolated according to a tissue-modified version of the standard Human Microbiome Project’s DNA isolation procedure. Genomic DNA from frozen lung tissue was extracted after tissue homogenization in Yeast Cell Lysis Buffer (EpiCentre) containing lysozyme (EpiCentre) by bead beating (TissueLyser II) with proteinase k (Invitrogen). DNA was purified with the Life Technologies PureLink kit according the manufacturer’s protocol (Invitrogen). A sterile water control (MoBio) was also processed along with all frozen tissue and used as background contamination control for DNA isolation, PCR and sequencing. Background contamination controls for tissue collection, pathology and sequencing were also collected through routine swabs after surgery and sequenced in conjunction with tissue samples. Specifically, the NCI-MD study tissues were isolated in a laminar flow hood to minimize contamination for downstream applications, using sterile forceps and gloves. Controls for contamination points during surgical tissue collection and pathological assessment included swabs from inside of the surgical tissue collection vessel before/after, pathology cutting board before/after, pathology knife blade before/after, gloves before/after, pathology ink bottle rim and collection tube for freezing before/after (See Additional file 1: Dataset 1). Briefly, swabs were dipped in Yeast cell Lysis buffer and area/object swabbed, then the swab was broken off into tube and frozen at -80. A negative control was also collected using 50 μL of MoBio PCR water as a mock sample (PCR_NC) and processed through DNA extraction with tissues to assess contamination from reagents, which was analyzed on three separate runs of MiSeq. The positive control was the High Even Mock Community (Broad Institute), which was also sequenced on three separate runs of MiSeq. The negative and positive control samples were spiked into four MiSeq runs at a similar concentration to that of the NCI-MD samples. To control for false grouping or batch affects, we randomized the tissue sample types (non-tumor (NT), tumor (T), immediate autopsy (ImA)) (with the exception of HB controls) across 5 separate sequencing runs of MiSeq (See Additional file 1: Dataset 2). The fifth plate consisted of duplicate samples and samples that had failed sequencing on previous runs of MiSeq.

Sequencing for the 16S rRNA gene was performed with 40 ng of sample DNA from 398 cases and 57 controls using primers for variable region V3-V5 with 16S rRNA gene sequence-specific portions based on Kozich et al. [58] with adapters for subsequent addition of standard Illumina dual indexes. PCR was performed using a Phusion DNA Polymerase High Fidelity kit (ThermoFisher). The cycling conditions were as follows: 98 ^°^C for 2 min, then 36 cycles of 98 ^°^C for 15 s, 60 ^°^C for 1 min 40 s, and 74 ^°^C for 1 min. PCR products were purified using the Agencourt AMPure XP kit according to the manufacturer’s instructions (Beckman Coulter). Second round PCR with Illumina dual-index oligos was performed using a Phusion DNA Polymerase High Fidelity kit (ThermoFisher) as following: 98 ^°^C for 2 min, then 6 cycles of 98 ^°^C for 15 s, 72 ^°^C for 20 s, and 72 ^°^C for 1 min. Samples were pooled, purified using Agencourt AMPure XP. Sequencing was conducted on Illumina MiSeq instrument using v3 600 cycles kit (see Additional file 1: Methods).

### Full-length 16S rDNA PCR reactions (PacBio)

Full-length 16S amplifications were performed using: 1µl of total DNA as template; 0.25 µM of the universal 16S primers F27 and R1492 with four different sets of asymmetric barcodes at (see Additional file 1: Table 7). and GoTaq Hot Start Master Mix (Promega) in a 50µl final volume. Cycling conditions were: 94 ^°^C, 3 min; 35 cycles of 94 ^°^C 30 sec, 54 ^°^C 30 sec, 72 ^°^C 2 min; following by a 5 min final elongation at 72 ^°^C. PCR products were cleaned with AxyPrep^™^ MagPCR (Corning Life Sciences) according to the manufacturer’s protocol and eluted in 40µl of water. Cleaned PCR products were quantified using the Bio-Rad QX200 droplet digital PCR (Bio-Rad) and QX200 EvaGreen® Supermix with primers F357 and R534 (see Additional file 1: Table 8) targeting the V3 variable region of 16S rDNA. Based on the results, amplicon libraries were normalized to the same concentration prior to pooling. Pooling was always performed using amplicon libraries with distinct barcodes. Multiplexing was performed with 2-4 libraries per pool.

### Pacific Biosciences circular consensus sequencing

Sequencing library construction was accomplished using the Pacific Biosciences (PacBio) SMRTbell^™^ Template Prep Kit V1 on the normalized pooled PCR products. Sequencing was performed using the PacBio RS II platform using protocol “Procedure & Checklist - 2 kb Template Preparation and Sequencing” (part number 001-143-835- 06). DNA Polymerase Binding Kit P6 V2 was used for sequencing primer annealing and polymerase binding. SMRTbell libraries were loaded onto SMRTcells V3 at a final concentration of 0.0125 nM using the MagBead kit, as determined using the PacBio Binding Calculator software. Internal Control Complex P6 was used for all reactions to monitor sequencing performance. DNA Sequencing Reagent V4 was used for sequencing on the PacBio RS II instrument, which included MagBead loading and stage start. Movie time was 3 hrs for all SMRTcells. PacBio sequencing runs were set up using RS Remote PacBio software and monitored using RS Dashboard software. Sequencing performance and basic statistics were collected using SMRT® Analysis Server v2.3.0. De-multiplexing and conversion to FastQ was accomplished using the Reads of Insert (ROI) protocol in the SMRT portal v2.3 software. Only reads with a minimum of 5 circular passes and a predicted accuracy of 90 (PacBio score) or better were used for further analysis. Each read was labelled in the header with the number of CCS (circular consensus sequence) passes and the sample designation using a custom ruby script, followed by concatenation of all reads into a single file for subsequent filtering and clustering.

### Filtering and OTU clustering of 16S rRNA gene sequence data

Initial screening for length and quality using QIIME (qiime.org) [59]. Reads containing more than five consecutive low quality base calls (Phred < Q20), were truncated at the beginning of the low quality region. Due to the low quality of the majority of R2 reads (Phred < Q20 and <150bp length), we used the R1 reads only for this analysis. Passing sequences were required to have high quality base calls (≥ Phred Q20) along a minimum of 75% of the read length to be included. After primer removal, final sequences containing ambiguous bases (Ns) or lengths less than 150bp were removed. High quality sequences were then screened for spurious PhiX contaminant using BLASTN with a word size of 16. Reads were then assessed for chimeras using USEARCH61 (de novo mode, 97% identity threshold for clustering). Non-chimeric sequences were screened for contaminant chloroplast and mitochondria using the RDP naïve Bayesian classifier, as well as non-specific human genome contaminant using Bowtie2 against the UCSC hg19 reference sequence. Finally, sequences were evaluated for residual contaminants using BLASTN searches of the GreenGenes database (v13.5). Filtered reads included those not matching any reference with at least 70% identity along 60% of their length. Exploratory assessment using BLASTN searches against the NCBI NT database indicated the majority unknown contaminant reads were amplified human genome sequence. High-quality passing sequences were subsequently clustered into operational taxonomic units using the open-reference operational taxonomic unit (OTU) picking methodology implemented within QIIME using default parameters and the GreenGenes database (99% OTUs) supplemented by reference sequences from the SILVA database (v111). Prior to downstream diversity analyses, the OTU table was rarefied to 5,500 sequences per sample. Prior to diversity analysis, contaminants were removed and again OTUs table rarified to 5,500 sequences per sample. Alpha diversity estimators and beta-diversity metrics were computed in QIIME with differential abundance analyses performed in R. In order to determine significant differences in beta diversity, we used the adonis function in the R package vegan to conduct PERMANOVA with Bray Curtis distance and 999 permutations. All sequences from MiSeq and PacBio data sets have been deposited at the following location: http://www.ncbi.nlm.nih.gov/bioproject/320383. See Additional file 1: Methods for details regarding PacBio sequence processing.

### TCGA RNA-seq data processing and alignment

In order to analyze all RNA-seq unmapped reads from TCGA lung cancer samples, we developed a custom metagenomic analysis pipeline using (i) MetaPhlAn2, (ii) Kraken, and (iii) Pathoscope [22]. First, all reads were filtered for quality using Trimmomatic (v0.32, minimum average quality > 20 over a 5bp sliding window, minimum final length^3^ 28bp) and searched for potential PhiX-174 contaminant using Bowtie2. Reads passing this filter were then mapped to the comprehensive NCBI Homo sapiens Annotation (Release 106) using Bowtie2 to remove any human-associated reads. The resulting non-human read set was then taxonomically assigned using (i) MetaPhlAn2, (ii) Kraken, and (iii) Pathoscope in parallel to evaluate consistency in the resulting profiles. Assignments from each method were aggregated at higher taxonomic levels (genus and species) for downstream statistical comparisons. The results from Pathoscope and its validation in other studies lead us to use these data for the remainder of the downstream analysis. Alpha diversity estimators and beta-diversity (Bray Curtis) metrics were computed in QIIME using genus and species level assignments with differential abundance analyses performed in R and Stata (v13).

### Statistical Analysis and Classification of Taxa Associated with Lung Cancer

Alpha and beta-diversity metrics were computed in QIIME with differential abundance analyses performed in R and Stata (v13). Mann-Whitney tests corrected for multiple testing (Benjamini–Hochberg (FDR)) were used to conduct initial comparisons between tissue type and histological subtype (AD or SCC) followed by multivariable logistic regression controlling for multiple confounders. A logistic regression model was constructed to estimate the odds of AD vs SCC for each taxa stratified by mutation status with and interaction term between the taxa and mutation added to the model. See See Additional file 1: Methods for details.

### TP53 gene sequencing and mutation analysis

Genomic DNA extracted from lung cancer tissues (n=107) was submitted for *TP53*-targeted sequencing using the MiSeq Illumina platform. For mutation analysis, 46 samples were SCC. The assay was targeted at the exons and proximal splice sites. Forward and reverse primers were tailed with Illumina Adapter tags for downstream next generation sequencing using the BioMark HD System (Fluidigm) and Access Array IFC chips and kits (Fluidigm). PCR products were indexed using an 8-mer oligo barcode. See Additional file 1: Table 3 lists sequences for primers used in the sequencing assay. Sequence results were processed and aligned to human genome and underwent QC requiring coverage > 100 reads with the variant (most SNVs had a read depth in the thousands) and minimum allele frequency > 10%. The 100-level cutoff for coverage allows to detect variations if the tumor fraction >~ 20% with 95% confidence, under the assumption of a diploid genome. The 10% allele frequency cutoff is derived from that same consideration. The variants called included all common polymorphisms. Because only tumor was sequenced, in order to score somatic mutations, those deemed to be germline were filtered out. These included SNVs present in dbSNP with high reported allele frequency (common polymorphisms). Also, SNVs in untranslated regions and introns were not considered, as their somatic status and functional implications are unclear. The presence of putative somatic exonic and splicing variants was corroborated in TCGA and COSMIC datasets. See Additional file 1: Table 2 for details.

### Fluorescent *in situ* hybridization analysis of *Acidovorax*

In order to confirm the presence *Acidovorax* in lung tumor tissue, fluorescently labeled probes were created for each bacterium. Genus or species-specific bacteria probes were hybridized using tumor tissues in addition to gram stain on each. Tumor tissues from cancer cases were fixed in OCT and sectioned frozen (10 μm). Prior to fixation in 4% paraformaldehyde sections were thawed at RT. Sections were washed in PBS and the probe (2 μL) was added to 90 μL FISH buffer (0.9 M NaCl, 0.02 M Tris pH 7.5, 0.01% SDS, 20% formamide). This solution was added to the section (20-100 μL) and placed in the hybridization chamber (46^°^C) for 3-18 hrs depending on probe used. Section were washed with twice (wash 1 - 0.9 M NaCl, 0.02 M Tris pH 7.5, 0.01% SDS, 20% formamide; wash 2 - 0.9 M NaCl, 0.02 M Tris pH 7.5, 0.01% SDS) and incubated at 48^°^C for 15 min. Slides were then dried for 10 min. Prior to visualization, DAPI and Vectashield was added slides. The probe used for FISH was: Acidovorax (CTT TCG CTC CGT TAT CCC, 5’ modification: Alexa Fluor 532). Representative fields were imaged using Zeiss 710 and a 100X objective for the probe. In addition to 2D images, Z stacks were also obtained for each bacterial probe and used to reconstruct 3D images and movies using Imaris software. Quantification of *Acidovorax* probe reactivity was conducted using 10 2D fields of two patients. At least 300 cells were counted per patient. Percentage (%) of cells with perinuclear probe reactivity was quantified using ImagePro Plus 6.0 software.

## Declarations

### Ethics approval and consent to participate

This study was approved by the Institutional Review Board at the NIH and all individuals participating in the NCI-MD case-control study signed informed consents for the collection of biospecimens, personal and medical information.

## Consent for publication

Not applicable.

## Availability of data and material

All de-identified data from this study will be made available upon reasonable request. Specifically, all sequencing data has been deposited under the bioproject number 320838 and is publically available at the following location: http://www.ncbi.nlm.nih.gov/bioproject/320383.

## Authors’ contributions

KLG, JAS and CCH conceived of the study, experiments, analyzed data, interpreted results and participated in writing and review; CD, SC, JO, AIR, VVB, MAP, SB, PSM, JEK, SVB, ASB, JPE, JCM, GDE, JAS and JRW sequenced samples and/or processed sequencing data and mutation analysis, and JRW, AJV and ECP conducted statistical analysis of data; AV, JAB, NM, TC, ANH, TMS, and MRW conducted FISH experiments and interpreted results; EDB and MAK provided assistance with procuring and processing biospecimens and clinical databases; BMR, AHD, and GT provided technical and data interpretation assistance and manuscript review. All authors read and approved the final manuscript.

## Acknowledgements

We thank the University of Marlyand Cancer Studies Team directed by Dean Mann, for their support, including Steven Schech for the collection of background swabs and specimens.

